# Comparison of sampling methods for small oxbow wetland fish communities

**DOI:** 10.1101/2022.03.10.483761

**Authors:** Dylan M. Osterhaus, Samuel S. Leberg, Clay L. Pierce, Timothy W. Stewart, Audrey McCombs

## Abstract

Throughout the world, wetlands have experienced degradation and declines in areal coverage. Fortunately, recognition of the value of wetlands has generated interest in preserving and restoring them. Post-restoration monitoring is necessary to analyze success or failure, thereby informing subsequent management decisions. Restoration of oxbow wetlands has become the focus of targeted restoration efforts to promote recovery of biodiversity and sensitive species, and to enhance ecosystem services. The fish communities of oxbows have been the subject of many monitoring studies. However, an optimal method for monitoring the fish communities of oxbows has not been described, thereby limiting our capacity to effectively manage these ecosystems. We compared four sampling methodologies (backpack electrofishing, fyke netting, minnow trapping, and seining) for fish community data collection with a primary objective of determining an optimal method for sampling fish communities in small oxbow wetlands. Seining and fyke netting were determined to be optimal methods for sampling oxbow fish communities. Backpack electrofishing and minnow trapping produced lower total catch and taxonomic richness values than seining and fyke netting. Although seining and fyke netting produced similar taxonomic diversity and abundance values, qualitative analysis revealed that seining caused greater habitat disturbance and potential stress to fish. Therefore, consideration must be given to how species present (especially sensitive species) within the wetland could be impacted by sampling disturbance when choosing between seining and fyke netting.

## Introduction

Wetlands are among the most impacted aquatic ecosystems, with declines in both areal coverage and habitat quality occurring as the global human population increases [1]. Fortunately, recognition of the value of wetlands has generated interest in preserving and restoring them [2]. Wetland restoration activities range in scope and complexity from watershed-scale undertakings to targeted fine-scale enhancements of habitats. As these activities have expanded, so has realization of the value of monitoring sites post-restoration to assess restoration successes or failures [3]. Identifying preferred methods of biological monitoring are imperative for comparing success of different restoration strategies and responses of different types of wetlands to these management actions [3].

Within the Midwestern United States, oxbow wetlands (hereafter oxbows) have recently become the focus of targeted restoration efforts due to the habitat that they provide for endangered Topeka Shiner *Notropis topeka*, and various other ecosystem services [4,5,6,7]. Oxbows are floodplain habitats consisting of river meanders that have been disconnected from the main channel through natural erosional processes, creating an off-channel lentic habitat [8,9]. These oxbow habitats are periodically reconnected with the main stream channel during high flow events which allows for colonization of oxbows by fish and dispersal of fishes from the oxbow back into the stream. As prairies, forests, and other forms of natural land cover in the region were replaced by cropland and pasture, streams were channelized, eliminating many oxbows [10,11]. As oxbows declined in number, their ecosystem services were also lost. In floodplain habitats, oxbows are especially valuable because they retain water, thereby reducing downstream flooding [5]. By sequestering nutrients, oxbows can contribute to improved stream water quality [5,6]. Additionally, oxbows contribute to increased biodiversity by providing essential wetland habitat for many species [6,12,13,14].

Restoration of oxbows has increased substantially over the last twenty years within the states of Iowa, Minnesota, and South Dakota. In Iowa alone, restoration of oxbows has increased from 1.2 restorations per year from 2001-2005 to 12.8 restorations per year from 2016-2020. Iowa oxbows are being restored because of ecosystem services described above, and because they appear to provide critically important habitat for the Topeka Shiner [4,5,6,7].

Accelerating oxbow restoration has led to increased monitoring of these ecosystems by The Nature Conservancy, the United States Fish and Wildlife Service, the United States Geological Survey and other entities. Monitoring projects are conducted in part to determine success or failure of restoring oxbow ecosystem services and habitat for Topeka Shiner. Monitoring typically involves collecting habitat data (depth, width, length, bank angles, substrate, canopy cover, etc.) and conducting plant and invertebrate community surveys. Given that oxbows are critical habitat for Topeka Shiner, research has also focused on fish communities of restored and unrestored oxbows [7,15,16,17].

While many studies have examined oxbow fish communities, an optimal sampling methodology for these habitats is not known, and some methods are likely more effective than others [15]. Additionally, given the differences in habitat morphology between oxbows and typical stream channels or wetlands, sampling methods that are more appropriate for streams or wetlands may not function as well when employed within oxbows. Sampling methodologies employed in previous oxbow fish community surveys include seining with a bag seine of variable sizes and numbers of seine passes [15], blocking the oxbow into four sections and sampling three while leaving one section as an undisturbed fish refuge [16], and “scoop” seining (1.2 x 1.2 m seine with 6.4 mm mesh)[18]. Because the Topeka Shiner is a rarely encountered species with a restricted geographic distribution and specific habitat requirements, it is important to identify an optimal fish community survey method for oxbows that are restored for Topeka Shiner habitat. An optimal methodology would produce consistently accurate information for species composition, taxonomic diversity, and abundance while minimizing stress to fish. Determination and implementation of an optimal sampling methodology for future studies will enable sampling without causing significant fish mortality, and efficient/effective sampling of the fish community.

In this study, we quantitatively compared values for three fish community metrics, total CPUE (catch per unit effort; number of fishes per 100 m^2^ oxbow surface area), species richness, and 10^th^-90^th^ quantile length ranges of fish collected that were obtained using four sampling methods. Additionally, we qualitatively compared these methods in terms of ease of use, stress to fish, habitat disturbance, ability to sample in dense vegetation, and ability to sample in deep water. The objective of our study is to compare performance of sampling gears for oxbow fish communities and given the focus of oxbow restoration on Topeka Shiner recovery, to make recommendations for appropriate sampling gears with consideration for sampling which may involve sensitive or endangered species.

## Materials and methods

Study sites consisted of 12 recently restored oxbows located in central Iowa (10 within the Boone River basin, two within the North Raccoon River basin). Oxbow length was measured along the centerline of the oxbow and ranged from 23 m to 139 m with an average of 77.4 m across sites. Oxbow width was measured as the wetted width at eight evenly spaced points along the length of the oxbow. These measurements were averaged at each site and average oxbow width ranged from 4.8 m to 20.4 m with an average of 10.0 m across sites. Depth was measured at three points along eight evenly spaced transects spanning the wetted width of the oxbow. These measurements were averaged at each site and average oxbow depth ranged from 0.56 m to 0.98 m with an average of 0.69 m across sites. Oxbow surface area was calculated by multiplying the average width of the oxbow by the length.

Fish communities at all oxbows were surveyed over a four-week period (18 May 2021 – 19 June 2021) using four sampling methodologies (backpack electrofishing, fyke netting, minnow trapping, seining). Each sampling methodology was used once at each oxbow for a total of 4 sampling events at each site. Seining is a frequently used fish sampling methodology in oxbows [7,15]. Fyke netting and minnow trapping are commonly used to sample fish in a variety of depressional wetlands [19,20,21,22]. Although infrequently used in wetlands, we included backpack electrofishing given the widespread use of this sampling method in fisheries field research.

Oxbow sampling sequence and order of methodology implementation were randomized (using a random number generator) to control for time effects on results. For each sampling method, all fish collected were identified to species and total numbers were counted. Total lengths of the first 50 recovered individuals for each species were measured. All fishes were subsequently released into the oxbow where they were captured.

### Sampling methods

Backpack electrofishing was performed by a single pass with a backpack electrofishing unit (Smith-Root LR-20B Backpack Electrofisher) across all wadeable habitat (< 1m in depth) by walking along the entire shoreline perimeter. One investigator carried the backpack, while two netters (one on each side of the backpack) collected fish. Electrofishing settings were determined based on conductivity of water in the oxbow (10-500 μs/cm, 200-300 volts; 500-800 μs/cm, 150-200 volts; 800-1000 μs/cm, 120-180 volts; >1000 μs/cm, 100-150 volts).

Fyke netting consisted of placing three, un-baited, mini-fyke nets (4.0 m lead, 0.6 cm mesh, largest hoop opening = 0.6 m×1.2 m) in each oxbow. Previous studies in depressional wetlands determined that 3 fyke nets were sufficient for determining relationships between fish community data, abundance and taxon richness of plants and invertebrates, and physical attributes of the wetland [22,23]. Nets were placed perpendicular to the shoreline within the open water zone at points spaced evenly along the length of the oxbow. Fyke nets were deployed for 24 h, and then retrieved.

Minnow trapping consisted of placing four, un-baited, minnow traps (2.5 cm opening, 0.6 cm mesh, 0.4 m long × 0.2 m diameter) at points evenly spaced along the length of the oxbow and at variable depths (0.3 m to 1.1 m) with two traps on each bank. Minnow traps were retrieved after 24 h of deployment.

Seining was conducted using a single pass of a bag seine (10.7 m×1.8 m, 0.6 cm mesh or 16.8 m×1.8 m, 0.6 cm mesh depending on oxbow width) along the entire length of the oxbow. For oxbows < 16 m in average width, the 10.7 m wide seine was used. For oxbows >16 m in average width, the 16.8 m wide seine was used. A single seine pass (rather than multiple passes) has been determined to be sufficient for surveying the fish community of an oxbow [7]. A single seine pass is more desirable compared to multiple seine passes given the reduced effort, and decreased fish stress and mortality [15].

### Fish community metrics and data analysis

To investigate effects of sampling method, we focused our quantitative analyses on three fish community metrics. Total CPUE was measured as the number of fish collected per 100 m^2^ of oxbow surface area. We chose to calculate CPUE in terms of total surface area, rather than amount of area sampled, as we sought to compare the performance of each gear type in terms of ability to sample the entire fish community within an oxbow. Also, given that the goal of sampling for our study was to document the entire fish community present within these closed systems at the time of sampling, we believe this to be the best way to compare sampling methods in terms of CPUE. Species richness was quantified as the number of species collected from one oxbow during one sampling event. The 10-90^th^ quantile range for lengths was calculated for each sampling event and these values were used for comparison. QQ-plots and histograms indicated that values for species richness and total CPUE were non-normal and heteroscedastic, therefore, these values were log transformed prior to analysis to meet normality and heteroscedasticity assumptions for associated analyses. Qualitative analyses focused on five metrics (ease of use, perceived stress to fish, perceived stress to habitat, ability to sample in dense vegetation, ability to sample deep water).

All analyses and figures were conducted and created using program R (version 3.6.3) [24] and packages **emmeans**, **ggridges**, **vegan**, **ggplot2**, **lme4**, and **plyr**. Mixed effects linear regression models were fit for each quantitative metric of interest (species richness, total CPUE, 10^th^-90^th^ quantile length range) with oxbow as a random effect and sampling method, and week included as additive predictors. A type III ANOVA was used to test for effects of sampling method on each fish community metric. The emmeans function was used to examine contrasts between sampling methods for each metric with results averaged over the levels of oxbow and week and P values adjusted using the Tukey method for comparing a family of four estimates. The critical value for statistically significant differences was P ≤ 0.05.

## Results

In total, 48 sampling events occurred (12 oxbows x 4 sampling events at each oxbow), resulting in collection of 32862 individual fish and 26 species (For a summary of fish catch data, refer to S1 Appendix). Within the sampling period, a fish-kill occurred at one oxbow, likely due to low dissolved oxygen levels as a result of decaying organic matter. Therefore, data from that oxbow were not included in analysis, resulting in 11 oxbows and 44 sampling events included in statistical analysis.

Fathead Minnow *Pimephales promelas* was collected from all oxbows and represented 73 % of the fish collected during sampling. Golden Shiner *Notemigonus crysoleucas* was the second most common fish sampled, representing nearly 9% of the total catch, although this species was collected from only 5 of the 11 oxbows. Seven of the 26 species collected represented over 97% of the total fish collected (S1 Appendix). Five species were collected from only one oxbow while two species were collected at each oxbow (S1 Appendix). Two Topeka Shiner were collected during sampling.

Sampling method had a significant effect on total CPUE (P < 0.001), species richness (P < 0.001), and fish lengths (P < 0.002; Fig 1). Total CPUE was significantly greater for fyke netting and seining than backpack electrofishing and minnow trapping (Fig 1; P < 0.001). Species richness was significantly greater for fyke netting, seining and backpack electrofishing than minnow trapping (Fig 1; P ≤ 0.009). Furthermore, across all sites, seining and fyke netting each collected 24 of the 26 total species encountered while electrofishing and minnow trapping collected only 19 and 7 species respectively (Fig 2). The range of fish lengths collected by seining, fyke netting, and backpack electrofishing were significantly greater than the range of fish lengths collected by minnow trapping (Fig 1; P < 0.04).

**Fig 1.**
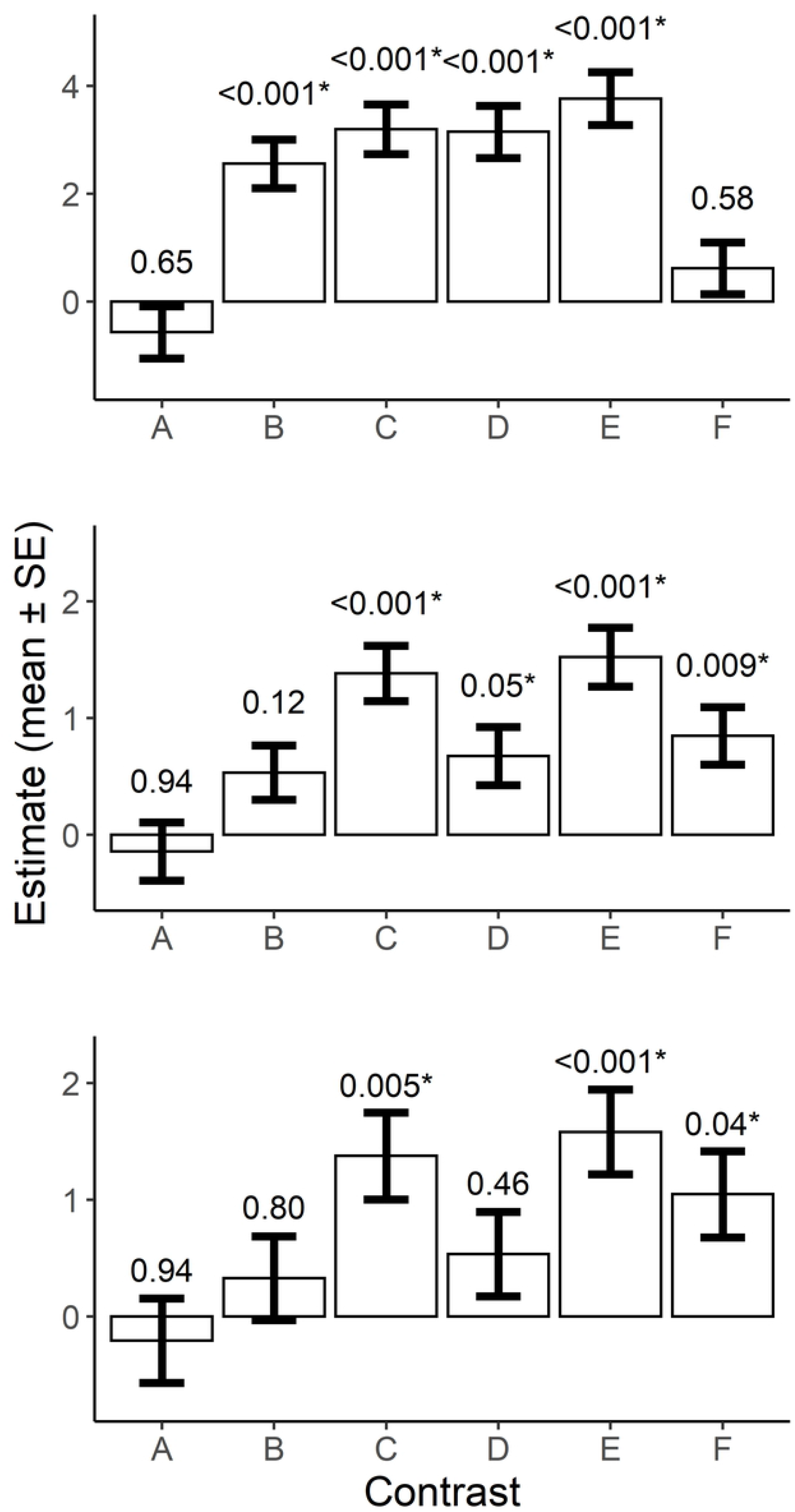
Pairwise-contrast statistics indicating extent of difference in CPUE (top panel), species richness (middle panel), and 10^th^-90^th^ quantile length ranges (bottom panel) as a function of sampling method. Contrasts include (A) fyke netting – seining (B) fyke netting – backpack electrofishing (C) fyke netting – minnow trapping (D) seining – backpack electrofishing (E) seining – minnow trapping (F) backpack electrofishing – minnow trapping. P values are based on Tukey’s adjustment for multiple comparisons. Asterisks denote a statistically significant difference.

**Fig 2.**
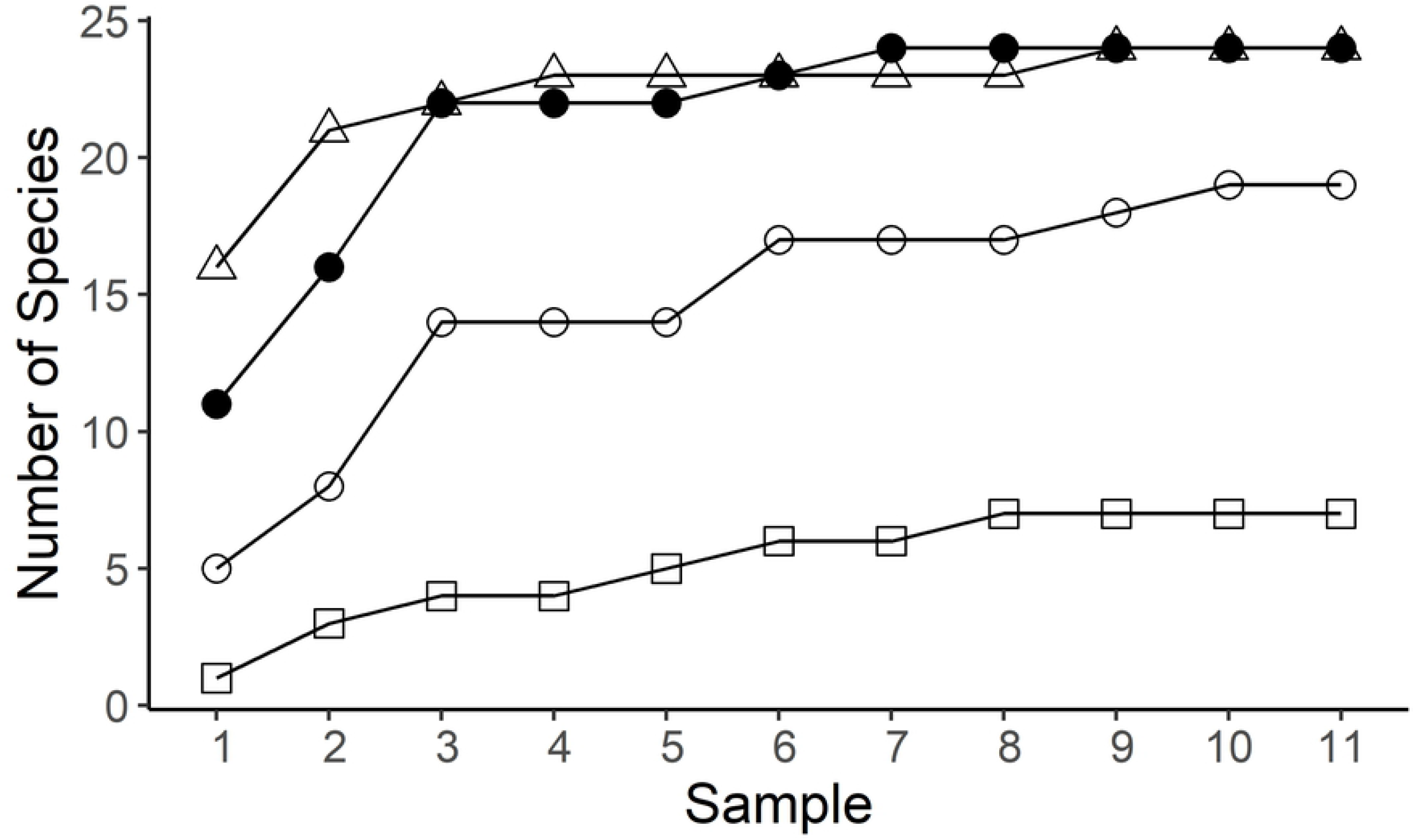
Species accumulation curves for each of the four sampling methods analyzed. Fyke netting represented with triangles, seining with closed circles, backpack electrofishing with open circles, and minnow trapping with squares.

### Backpack electrofishing

Total CPUE values obtained via backpack electrofishing were low compared to other methods, indicating that this method does not perform well in sampling oxbow fish communities (Fig 1). Species richness values produced by backpack electrofishing (range = 1–11 species per oxbow based on 11 sampling events) were lower than fyke netting (2–16 species) and seining (2–16 species), indicating that backpack electrofishing fails to detect many species occupying an oxbow (Table 1). Backpack electrofishing was successful in collecting the 13 most common of the 26 total species collected (S1 Appendix). However, seven of the 13 least common species were absent from collections made via backpack electrofishing. Length frequency distribution data indicate that backpack electrofishing samples fish similarly to fyke netting and seining and collects a larger range of fishes than does minnow trapping (Figs 1, 3).

**Figure 3.**
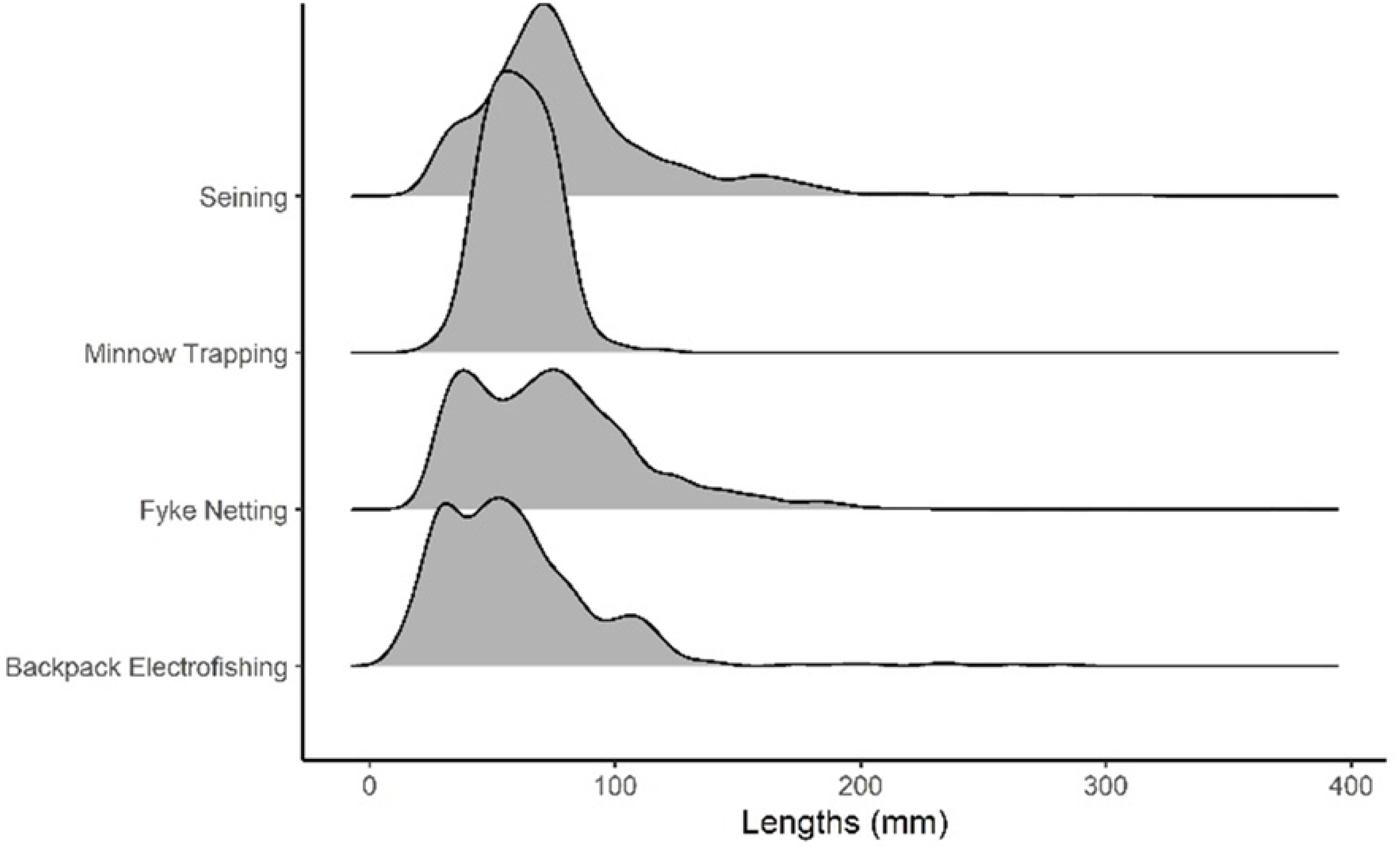
Length frequency distribution for lengths as sampled by each sampling method (n = 1716, 215, 1344, and 491 respectively for seining, minnow trapping, fyke netting, and backpack electrofishing. Height of the shaded region for each sampling method along y-axis represents the frequency of fishes of that length

**Table 1.**
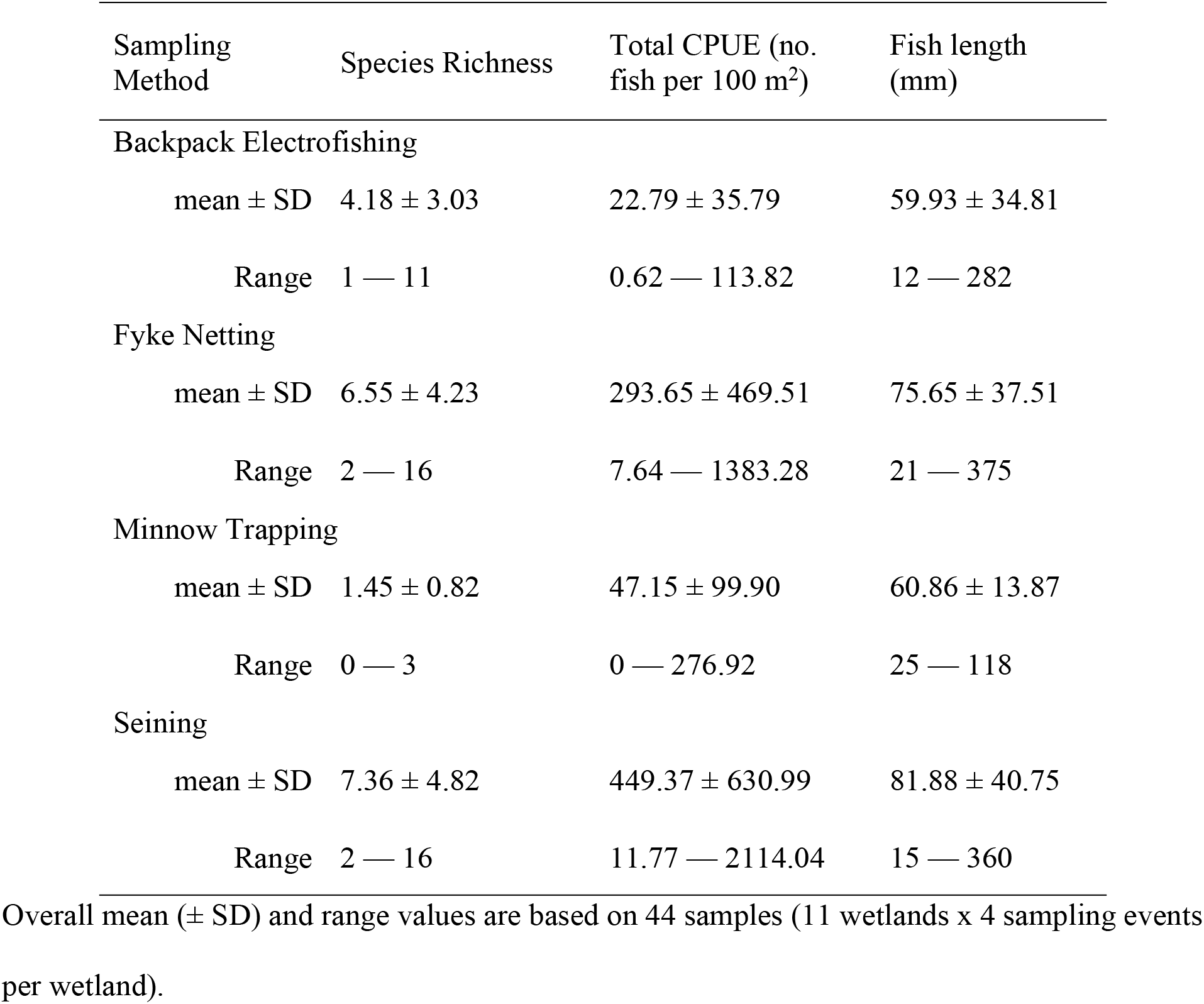
Descriptive statistics by sampling method for each quantitative metric.

### Fyke netting

Total CPUE values from fyke netting were comparatively high, indicating that this technique performs well in capturing fishes occupying the oxbow (Fig 1). Furthermore, fyke netting resulted in the collection of 24 of the 26 fish species recorded during the study, missing only Rock Bass (representing the lowest proportion of total catch, collected at one oxbow via seining) and Yellow Perch (representing the fourth lowest proportion of total catch, collected at one oxbow via seining) (S1 Appendix; Fig 2). Additionally, a single Topeka Shiner was collected via fyke netting at one oxbow. These findings, in combination with the high species richness values, indicate that fyke netting is effective at detecting rarer species. Length frequency distribution results from fyke netting indicate that this method samples a greater range of fish sizes present in the oxbow compared to other methods used (Figs 1, 3).

### Minnow trapping

Total CPUE values from minnow trapping were the lowest of the four methods tested (Fig 1). Minnow trapping resulted in collection of seven species in total, but no more than three species were collected during any one sampling event (Fig 2; Table 1). Length ranges of fish collected via minnow trapping were significantly smaller than those collected via backpack electrofishing, fyke netting, and seining (Fig 1). Minnow trapping appears to be more size-selective for fish in the length range of 40 to 90 mm, in comparison to other sampling methods (Figs 1, 3).

### Seining

Total CPUE values indicate that sampling performance is high for seining. Seining resulted in collection of 24 of the 26 species found during this study, missing only Black Crappie (tenth lowest proportion of total catch, collected at one oxbow via fyke netting) and Carmine Shiner (third lowest proportion of total catch, collected at one oxbow via fyke netting; Fig 2; S1 Appendix). The range of species richness values generated by seining (2-16 species) was identical to that of fyke netting (Table 1). Furthermore, a single Topeka Shiner was collected via seining at one oxbow. As was true for fyke netting, it appears that seining effectively samples rarer species of fish. Seining does not appear to be size-selective (Figs 1, 3).

## Discussion

### Comparison of sampling methods for fish community sampling

Our findings indicate that seining and fyke netting are comparable in how they sample oxbow fish communities and perform better than the other sampling methodologies evaluated in this study. These conclusions are based on high total CPUE and species richness values, unbiased sampling of various size classes, and ability to sample deep-water habitat. While fyke netting and seining both appear to perform well in sampling oxbow fish communities, there are differences in ease of implementation of each gear type as well as the level of habitat disturbance generated.

It is possible that under sampling of the fish community by fyke netting and minnow trapping occurred as the number of nets and traps was not scaled to oxbow size and our findings should be considered with this in mind. However, other studies have found three fyke nets to be sufficient for determining strong relationships between fish community data, abundance and taxon richness of plants and invertebrates, and physical attributes of depressional wetlands that were larger in size than our oxbows [22,23]. Furthermore, our findings indicate that fyke netting with four nets regardless of oxbow size performed better than backpack electrofishing and minnow trapping and as well as seining in terms of species richness values, and total CPUE. Also, minnow trapping is likely an ineffective gear type for sampling oxbow fish communities given the size selectivity of the gear, and previous studies have reached this same conclusion [25, 26,27].

### Backpack electrofishing

Backpack electrofishing was a difficult technique to use in oxbows. First and foremost, turbidity is typically high within oxbows and sediment is easily suspended. Given that efficacy of backpack electrofishing relies heavily on the netter’s ability to see stunned fish [28], relying on this sampling method in turbid systems is not recommended. Furthermore, the sampled oxbows frequently contained areas in which water depth was > 1m. To maintain the backpack electrofishing unit’s functionality by avoiding the submersion of the circuitry and power source, much deep-water habitat was unsampled. Presence of dense submerged plants also created difficulty because fish stunned by the electrical currents became ensnared in vegetation and did not surface.

Backpack electrofishing can also be stressful to fish [29,30]. Handling stress may be high as this method does not lend itself well to holding fish for data collection after sampling. Fish are transferred from the collection net to a larger holding container within the oxbow, which at times involves a significant amount of handling. In terms of disturbance to the habitat, backpack electrofishing in our study involved three individuals walking through the entire wadeable portion of the oxbow. This leads to heavy disturbance of sediment, increased turbidity, and displacement of vegetation throughout the entirety of the oxbows wadeable habitat.

### Fyke netting

Fyke netting had multiple drawbacks (amount and weight of gear, time requirements) and benefits (low stress to fish, ability to sample in deeper water and dense vegetation). The amount and weight of gear needed for fyke netting created difficulty when hauling gear to study sites (in our case, a distance up to 1 km). Fyke netting is time-consuming as nets are typically deployed for 24 h, which necessitates site visits on successive days. Furthermore, habitat disruption by fyke netting was seen to be not as extensive as that caused by more active forms of sampling, including backpack electrofishing and seining in this study. For fyke netting, habitat disruption within an oxbow is generally confined to placement locations of fyke nets and does not result in widespread disturbance of vegetation and benthic substrate.

During fyke netting, fish can be quickly transferred from the net to a holding container and then counted and released immediately which limits handling time. However, fish are restrained within the net from the time of their entry until when the nets are processed. This restraint may disrupt normal behavioral patterns and expose fish to increased predation risk if the net is occupied by piscivores such as Green Sunfish, Largemouth Bass, or turtles. Handling stress from fyke netting is likely similar to that caused by minnow trapping but lower than seining and backpack electrofishing. Additionally, few mortalities were noted during the use of fyke netting for this study.

### Minnow trapping

Minnow trapping is easy to implement as a sampling practice given the small size and low cost of traps and the ease with which they can be placed into the oxbow. However, minnow trapping is likely an ineffective sampling methodology for oxbows with significant aquatic vegetation. Aquatic macrophytes along the bottoms of an oxbow may inhibit effectiveness of minnow traps because entry funnels may become clogged or covered with vegetation. Also, given the small size of the funnel leading to the trap entrance, it is likely that each trap is sampling only a small proportion of the total habitat present at an oxbow in comparison to backpack electrofishing, fyke netting, and seining. To sample the entirety of the oxbow, a larger number of minnow traps is likely needed. However, even if enough minnow traps were used to adequately sample the entire oxbow, the size selectiveness of minnow traps represents a significant drawback in terms of the use of this gear.

Minnow trapping causes minimal disturbance to oxbow habitat as traps can be deployed from the shoreline, without an investigator having to enter aquatic habitat. While habitat disturbance may be limited, stress to fish collected via minnow trapping may occur. Similar to fyke netting, fish confined within a minnow trap may experience disruption to normal behavioral processes, such as feeding, for up to 24 h. However, unlike fyke netting, given the size selectiveness of minnow trapping, fish in traps are unlikely to experience increased predation risk because larger piscivores are unable to enter traps. Overall, few mortalities were noted during the implementation of minnow trapping during this study.

### Seining

Seining can be effective for sampling deep water up to the height of the seine (1.8 m), as investigators pulling the seine may remain along the shallow shoreline of the oxbow during sampling. Documented detection probabilities range from 40% to over 90% for fish species in similar oxbows using similar seining techniques [16]. However, for oxbows with dense aquatic vegetation or an abundance of woody debris/boulders, seining becomes arduous given the active nature of this method. Additionally, in oxbows with aquatic vegetation, seining can lead to significant habitat disturbance within the oxbow as the lead line of a seine may disturb an oxbow’s floor and up-root aquatic vegetation. Of the sampling methods employed by this study, seining likely generated the highest levels of habitat disturbance.

During seining, fish are held entirely in the water until the moment each individual is counted, which may reduce handling stress. Also, the amount of time in which an individual fish is impacted is minimal compared to passive forms of sampling which may result in a fish being trapped for 24 h. However, when oxbows with significant aquatic vegetation are sampled, a risk exists for entanglement of fish within aquatic vegetation collected by the seine. High mortality of fishes collected in seines can occur under these conditions.

## Conclusions

Minnow trapping is seemingly ineffective in accurately describing the oxbow fish community using the methods we employed. Other investigators determined that the catch of minnow traps is biased towards common species, is size-selective, and performs poorly in comparison to seining; our findings corroborate these studies [25,26,27]. Given the inability to sample the fish community effectively and unbiasedly (as determined by our study and others), we do not recommend use of minnow trapping as a sampling method for oxbow fish communities. Furthermore, given the low CPUE values, ineffectiveness in sampling rarer species, poor performance in comparison to fyke netting, difficulties in sampling deep and vegetated water, and high disturbance to habitat, we do not recommend backpack electrofishing as a sampling methodology for oxbow fish communities. However, we do recommend both seining and fyke netting as optimal methods for oxbow fish community sampling. Quantitively, fyke netting and seining appear to be comparable in fish community sampling performance. In our study, we found differences in the ranges of lengths of fish collected, however, both sampling methods appear to sample sizes classes without bias (Fig 1).

When choosing between fyke netting and seining, consideration must be given to the amount of time needed to conduct sampling, sensitivity of potentially encountered species to habitat disturbance and handling, habitat characteristics (such as aquatic vegetation), accessibility of the oxbow, and time of year. For studies constrained by a short timeline, seining may be more desirable as an oxbow can be visited once and sampled via seine within a few hours, while fyke netting may require two days and two visits to the oxbow to complete. For oxbows containing large quantities of aquatic vegetation or woody debris or boulders, fyke netting is the preferred sampling method. However, given the weight and difficulty in transporting fyke nets from a vehicle to the oxbow, seining may be preferable for oxbows that are not readily accessible by vehicle.

In addition to sampling method performance, consideration must be given to potential stress experienced by fish (either directly via handling or indirectly via habitat disturbance) as surveys are conducted. These considerations are especially significant when handling species of greatest conservation concern. For studies sampling oxbows which may contain species sensitive to habitat disturbance, fyke netting may be preferable to seining given the decreased habitat disturbance of this gear and special considerations should be given to timing of the spawning season for the species. For example, if sampling occurs during the breeding season for nest building centrarchid sunfishes such as the Green Sunfish, and Bluegill (Topeka Shiner nest in association with each of these species) [18,31,32,33], this reproductive activity could be compromised if the nests of the centrarchids are destroyed by seining. Therefore, fyke netting is recommended as the preferred sampling method during the Topeka Shiner breeding season.

In summary, the monitoring of oxbows, both before and after restoration, is necessary to understand impacts of restoration on the fish community present within the oxbow. We must understand how restoration impacts the fish community to improve management practices that maximize benefits for target species. Based on results from deploying 3 mini-fyke nets for 24 hours in wetlands ranging from 130 m^2^ to 2200 m^2^ in surface area, and results obtained by seining the entire wetland area, we determined that fyke netting and seining performed equally well in sampling oxbow fish communities. However, we conclude that fyke netting is preferable if the wetland is suspected to contain species sensitive to handling or habitat disturbance.

## Acknowledgements

We thank our field technicians, S. Grinstead and C. Wood, for the many hours spent in the field collecting the data presented within this manuscript. We also thank the various collaborating entities and individuals who have made this project possible, either through funding or support of the research efforts. Collaborators for this project include C. McKinney, K. Wilke, A. Kenney, J. Olson, D. Weissenfluh, Iowa Soybean Association, Syngenta Crop Protection LLC, The Nature Conservancy, and the U.S. Fish and Wildlife Service. Finally, we thank the many landowners whose support was vital for the success of this project.

## Supporting information

S1 Appendix— Abundance values for fish species captured in this study, including number of individuals in total and by sampling method, and its overall relative abundance (percent of total individuals). Values are based on 11 sampling events for each gear type in each wetland (18 May – 11 June).

